# denim: an R package for deterministic compartmental models with flexible dwell time distributions

**DOI:** 10.1101/2025.08.05.668369

**Authors:** Thinh P. Ong, Anh T. Q. Phan, Lam M. Ha, Marc Choisy

**Affiliations:** Oxford University Clinical Research Unit, Ho Chi Minh city, Vietnam; Centre for Tropical Medicine and Global Health, Nuffield Department of Medicine, University of Oxford, Oxford, United Kingdom

**Keywords:** Compartmental model, sojourn time distribution, dwell time distribution, R package, semi-Markovian, competitive risks

## Abstract

1. Compartmental models are widely used for dynamical systems where states are discrete such as in infectious diseases epidemiology with the so-called SIR or Susceptible-Infectious-Recovered framework. For mathematical simplicity, rates of transition between compartments are generally assumed to be independent of the dwell time (or secondary time scale in survival analysis): they are either constant or dependent on the epidemiological time (or primary time scale in survival analysis) only, either directly (e.g. environmental or behavioral forcings in epidemiological models) or indirectly through dependence on other variables of the system (e.g. the force of infection in epidemiological models). In some domains of application, this memoryless assumption leads to distributions of dwelling times that are incompatible with those observed on data which can lead to serious problems since the model predictions are highly sensitive on the exact shape of these distributions.
2. Here we propose a deterministic, continuous-variable, numerical modelling approach that allows full flexibility on the dwell time distributions. The accompanying denim package provides a user-friendly interface to implement our proposed method through a dedicated language for model definition.
3. The package is open-source and available on CRAN. With more detailed data on the clinical process of infections becoming available, the denim package will be extremely useful for building more realistic epidemiological models providing more accurate projections.

The denim package is publicly available for download on CRAN (https://cran.r-project.org/web/packages/denim/index.html) and GitHub (https://github.com/thinhong/denim). Additional documentation can be found on the denim webpage (https://drthinhong.com/denim/). Bug reports can be submitted via Github Issue.

## 1. Introduction

Compartmental models are widely used in various domains of applications dealing with the dynamics between discrete states. This is the case for example in epidemiology, where modelers typically divide the population into susceptible, infectious, and recovered with specified transition rates between them. In their classical and most used form, these models are Markovian (i.e., memoryless), meaning that the rate at which a compartment is exited is independent of the duration spent in the compartment (or the secondary time scale in survival analysis). The main motivation of this assumption is tractability where these models can easily be implemented using simple ordinary differential equations (ODEs) (Hong et al., 2024a). Furthermore, under the Markovian assumption, modeling transitions from one compartment to multiple others as competing risks is straightforward, as the total exit rate from the compartment is simply the sum of the individual transition rates. It also allows to express deterministic systems of ODEs as discrete counting processes, which can be easily simulated and parameterised (eg. via Gillespie algorithm) in stochastic implementations (Zachreson et al., 2022).

The consequence of the Markovian assumption however is that it implicitly involves a specific distribution of dwelling time that can be incompatible with observed data. For example, constant rates of transitions between the latent/infectious and recovered compartments in an epidemiological model implies distribution of latency/infectious durations that are exponentially distributed with both large variance and highest densities of probability for the shortest durations - a feature that is not supported by the empiric distributions of these durations which are typically normally distributed with small variance around a central tendency (NISHIURA & EICHNER, 2006; Wearing et al., 2005). Theoretical analyses further show that the shape of the exact distributions of dwell times greatly affect the model’s trajectory (Keeling & Grenfell, 1998; Kenah & Miller, 2011; Krylova & Earn, 2013; Lloyd, 2001). Wearing et al. (2005) illustrated that assuming exponential infectious periods always underestimate the basic reproductive ratio.

An easy relaxation of this strong assumption of exponential distribution proposed in the literature is to assume that dwell times follow an Erlang distribution. Indeed, since an Erlang distribution of rate *λ* and shape k is the sum of k independent exponential distributions of rate *λ*, it can easily be implemented with ODEs through the use of the linear chain trick (LCT) (Hurtado & Kirosingh, 2019). Even though this 2-parameter distribution adds flexibility to the distribution of dwell times, there are still many instances when it does not provide a great fit to data. Furthermore, the computer code for the LCT typically grows with the number of sub-compartments in the chain which renders the code development and maintenance cumbersome. Recent studies proposed an alternative approach to incorporate any type of parametric distributions by formulating the compartmental model as a system of delay integro-differential equations (Hernández et al., 2021; Hong et al., 2024b) then solving it numerically. A separate work by Tallis (1994) also discusses how competing risks can be incorporated into compartmental models that are under the non-Markovian assumptions. However, to our knowledge, no prior work has introduced a simulation algorithm capable of handling compartmental models with arbitrarily distributed competing risks. Additionally, in terms of implementation, no existing tools offer a simple way to model arbitrary dwell time distributions. To do so usually requires tedious and time-consuming work around, compromising the code readability.

In this paper, we provide a semi-Markovian modelling framework implemented in the R package *denim* that allow users to specify any dwell time distribution, including non-parametric forms and direct integration of empirical data, that will allow more realistic modelling of progression and delay. Model structures are easy to define and highly customisable with support for competing risks. The syntax is intuitive and designed for seamless integration with existing workflows using *deSolve (Soetaert et al*., *2008)* or *odin (FitzJohn, 2019)*, enabling users to adapt their current models with minimal effort.

## 2. Modeling approach

Our framework considers a deterministic, discrete-time, and continuous variable compartmental model in which all the compartments are further subdivided into sub-compartments, the values of interest being transition probabilities between sub-compartments.

To demonstrate denim’s algorithm, consider a basic compartment model with 3 compartments *X, Y, Z*. Individuals in compartment *Y* comes from *X*, and will move to *Z*. Consider that the time individuals stay in *Y* before moving to *Z* follows a discrete probability distribution *P*_*y*_ = *p*_1_, *p*_2_,…., *p*_*n*_ with 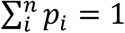 where *n* is the maximum number of time steps individuals can stay in compartment *Y*. Compartment *Y* is then further split into sub-compartments, where sub-compartment *Y*_*i*_ represents the number of individuals that has been in the *Y* compartment for *i* time steps.

**Figure 1:**
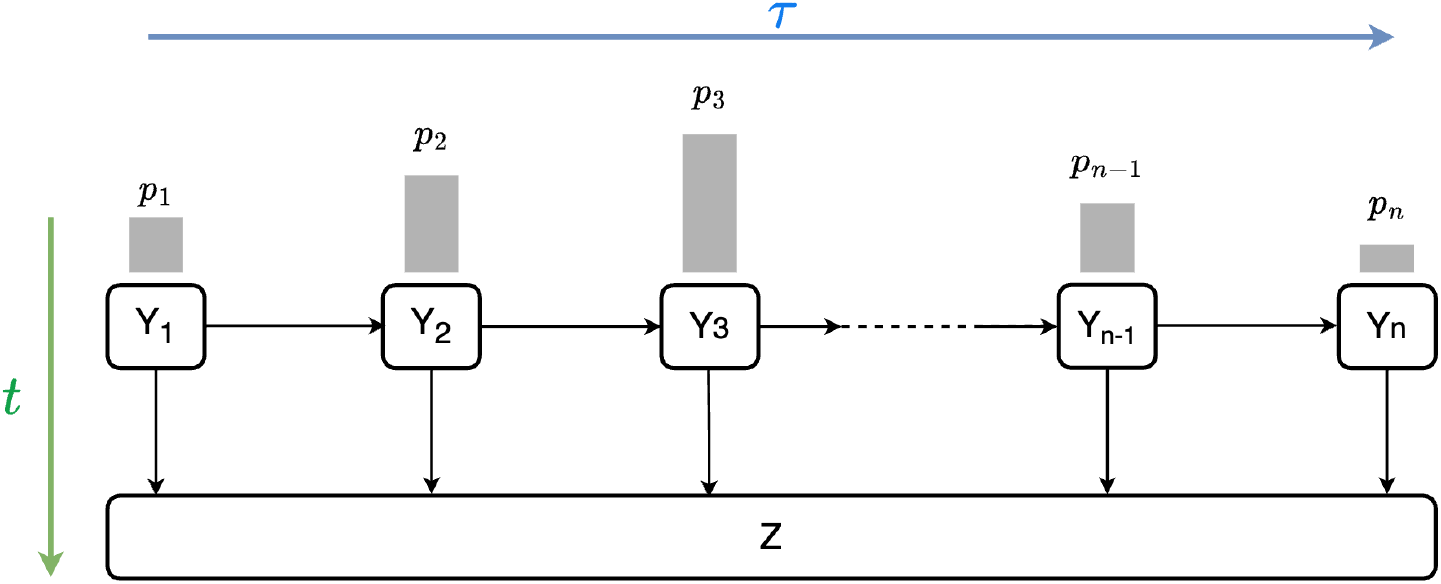
Visualization of the sub-compartments in denim. p_i_ represents the proportion of individuals in Y that transition to Z from each sub-compartment Y_i_. There are 2 timelines presented, t represents the primary time scale (i.e., calendar time) while τ represents the secondary time scale (i.e., dwell time, or time since entering compartment Y).

At each time step, a proportion *q*_*i*_ of individuals in *Y*_*i*_ (where *i* < *n*) move to *Z* compartment, while the remaining 1 − *q*_*i*_ move to *Y*_*i*+1_. The central problem in this sub-compartment approach is to determine how the transition proportion *q*_*i*_ relates to the target dwell time distribution *p*_*i*_.

To derive that relationship, we first reformulate our problem under the framework of survival analysis where the event of interest is “transitioning from *Y* to *X*”. The relationship between compartmental model and survival analysis has been explored in previous studies (Hay et al., 2024; Tallis, 1994) and is discussed in greater details in Supplementary Material 1. Under the survival analysis framework, the distribution *P*_*y*_ = *p*_1_, *p*_2_,…., *p*_*n*_ can be interpreted as a discrete estimation of a probability density function *f*(*τ*) of dwelling times, such that:

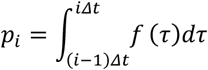

Where *Δt* is the duration of each time step.

*q*_*i*_ (probability of leaving *Y*, given that individuals have stayed for *i* time steps i.e., P(Z|*Y*_*i*_)) is thus an estimation for hazard rate *h*(*τ*) in survival analysis. The relationship between *h*(*τ*) and *f*(*τ*) is represented by the following formula.

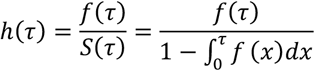

In discrete time, *q*_*i*_ can then be estimated using *p*_*i*_ as followed.

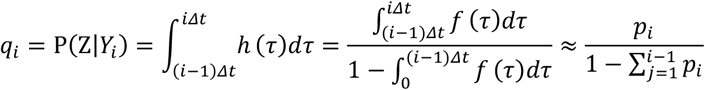

A visualization of transitions between sub-compartment *Y*_*i*_ and *Z* is provided below.

**Figure 2:**
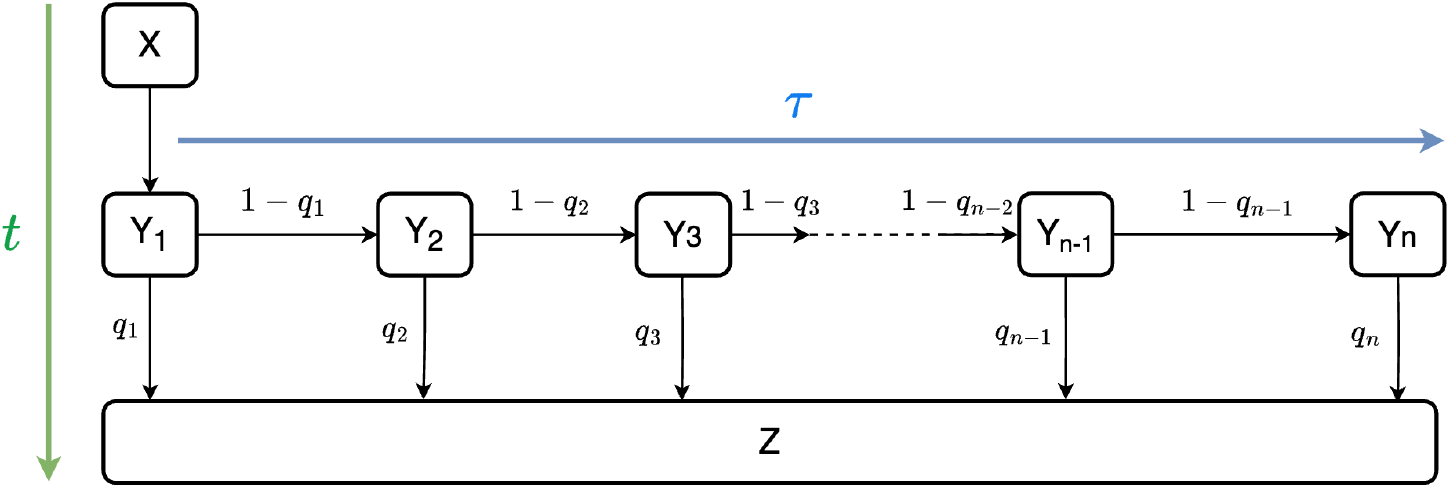
Visualization of the transition proportions of each sub-compartment Y_i_

This sub-compartmental structure can be used as a numerical method to solve the following system of delay integro-differential equations.

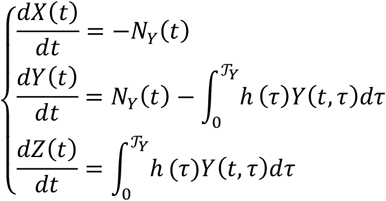

Where:

- Where *N*_*Y*_ (*t*) is the population of *X* that transition to *Y* at time *t*.
- 𝒯_*Y*_ denotes the maximal dwell time in *Y* compartment.
- *Y*(*t, τ*) is the sub-population of *Y* at time *t* that have been in *Y* for a duration *τ* (i.e., *Y*(*t, τ*) = *S*(*τ*)*N*_*Y*_ (*t* − *τ*)). The sub-compartment *Y*_*i*_ can thus be interpreted as 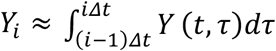

When using the package denim, users only need to define a set of dwell time distributions, and the package will automatically compute all transition probabilities *q*_*i*_ using the formulation above. The dwell time distribution could be either one of denim’s built-in parametric distributions or a vector of values corresponding to the ordered values of a histogram with a bin width equal to the time step of the model. Currently supported parametric distributions include: exponential, gamma, log-normal, weibull.

With the transition probability from each sub-compartment computed, the simulation can be carried out by updating at each time step the value of each compartment from incoming and outgoing individuals. In the given example, the out population for *Y* (or the in-coming population for *Z*) is 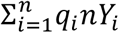 where *nY*_*i*_ is the population that have stayed in compartment *Y* for *i* time steps (i.e., population of *Y*_*i*_ sub-compartment).

### 2.1. Transition to multiple outgoing compartments

To model transitions from one to multiple compartments, there are 2 ways to define the transitions: (i) as multinomial transitions, or (ii) as competing risks depending on what information modelers currently have.

Consider a scenario where individuals in compartment *Y* can transition to an additional compartment *V*. The information we may have includes:

1. Distribution of the time individuals stay in Y, denoted *f*_*Y*_ (*τ*).
2. Distributions of the time individuals transitioning from Y to Z and from Y to V stay in Y (denoted *f*_*Y*→*Z*_(*τ*) and *f*_*Y*→*V*_(*τ*) respectively).
3. Proportion of individuals in Y compartment that end up in Z and V, denoted *w*_1_ and *w*_2_ respectively.

If information 2 is available (*Scenario 1*), transitions can be modeled using competing risks framework. Alternatively, if we have information 2+3 (*Scenario 2*) or 1+3 (*Scenario 3*), the transitions can be modeled using multinomial framework. In other words, multinomial is used when we want to specify how individuals are distributed across *k* downstream compartments following a proportion *w*_*m*_ where 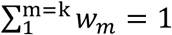. A visualization of all the scenarios is provided below.

**Figure 3:**
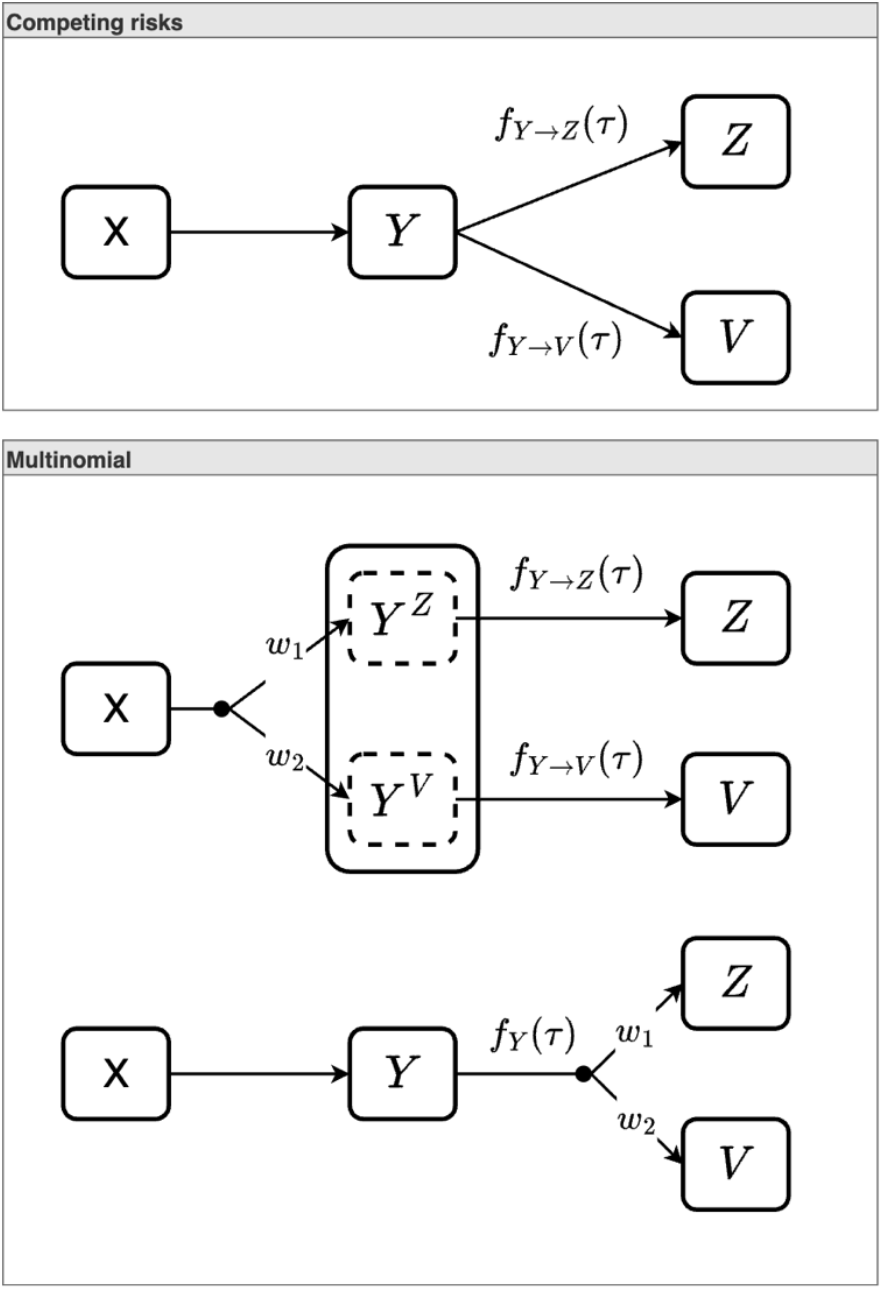
Visualization of the 3 scenarios. The lines extending from the black dot indicate the proportion of individuals that will eventually enter each outgoing compartment at equilibrium.

Multinomial transition is implemented by having *k* sub-compartment chains corresponding to *k* outgoing compartments. Upon entry, a proportion *w*_*m*_ of the population will be allocated to sub-compartment chain *m*. For example, to implement an addition compartment *V* where 80% of *Y* goes to *V* and the remaining 20% goes to *Z*, we do the following: (i) create 2 sub-compartment chains for *Y* → *V* and *Y* → *Z* transitions, then (ii) when a population of *n* enters *Y*, 0.8 * *n* goes to *Y* → *V* chain and 0.2 * *n* goes to *Y* → *Z* chain and (iii) the outgoing population to *V* and *Z* are computed from these sub-compartment chains independently. Under *Scenario 2*, out-going population for each chain computed using *f*_*Y*→*Z*_(*τ*) and *f*_*Y*→*V*_(*τ*). To implement *Scenario 3*, it is equivalent to setting *f*_*Y*→*Z*_(*τ*) = *f*_*Y*→*V*_(*τ*) = *f*_*Y*_ (*τ*), a proof for this is provided in Supplementary Materials 1.

**Figure 4:**
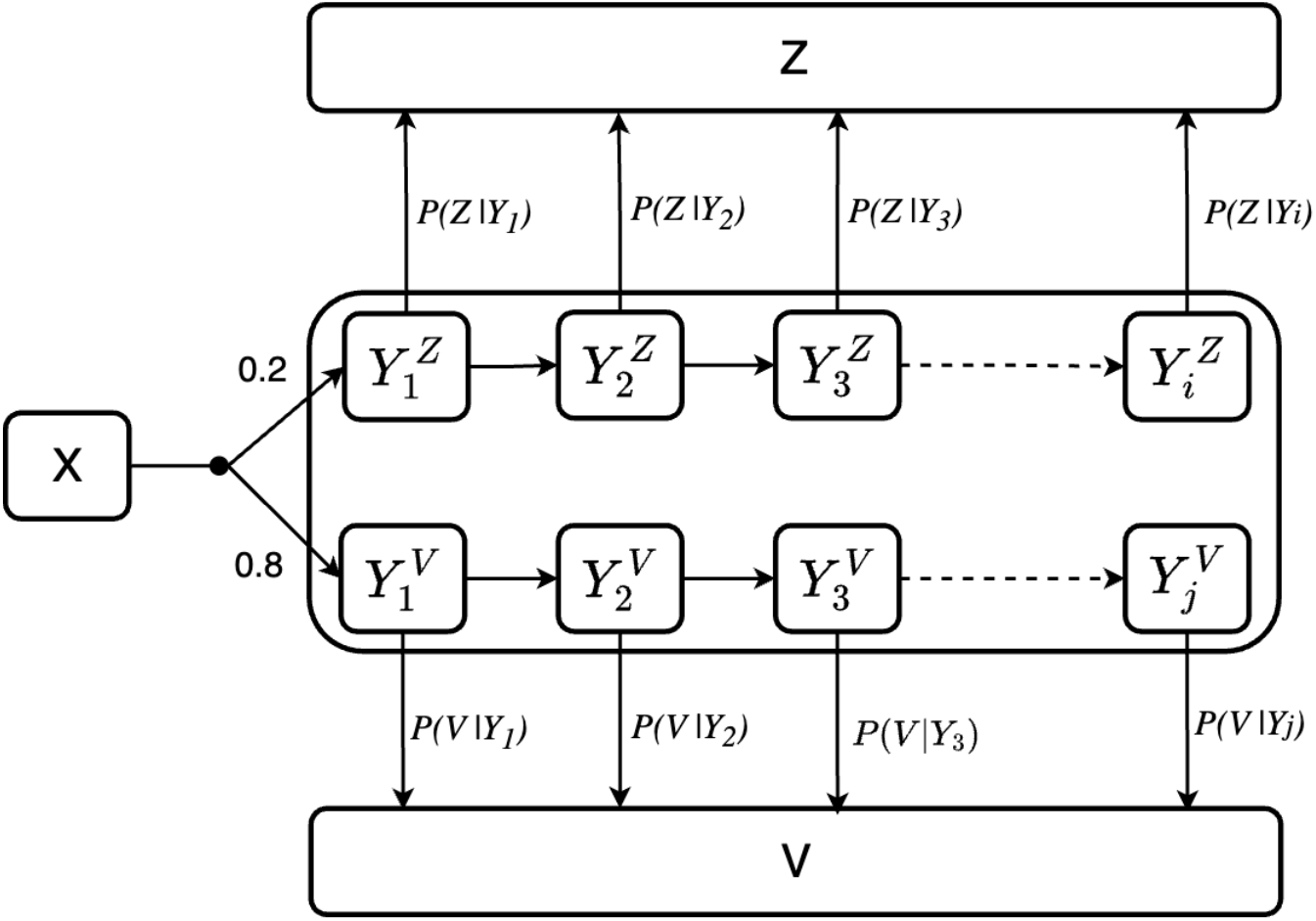
Visualization of the sub-compartment chains in multinomial. Y^Z^and Y^V^denote the sub-populations of Y that will transition to Z and V respectively. The total population that goes to Z is computed by 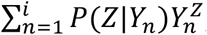, and population that goes to V is 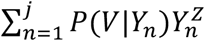.

Outgoing transitions are modeled as competing risks when we have the distributions of transitioning time to each out-going compartment, but without the knowledge of *p*_*m*_. This is implemented by using only one sub-compartments chain, the length of which is determined by the maximum waiting time amongst all outgoing transitions. The outgoing population is then computed from this one sub-compartments chain. For example, suppose *Y* → *V* and *Y* → *Z* are competing risks, with maximal waiting times *n* and *m* respectively, where *n>m*. To model this, do the following: (i) create a sub-compartment chain with length *n*, (ii) compute the population that goes to *V* as 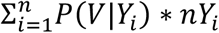 and that of *Z* is 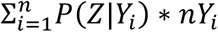 where *P*(*Z*|*Y*_*i*_) = 0 when *i*>*m*.

**Figure 5.**
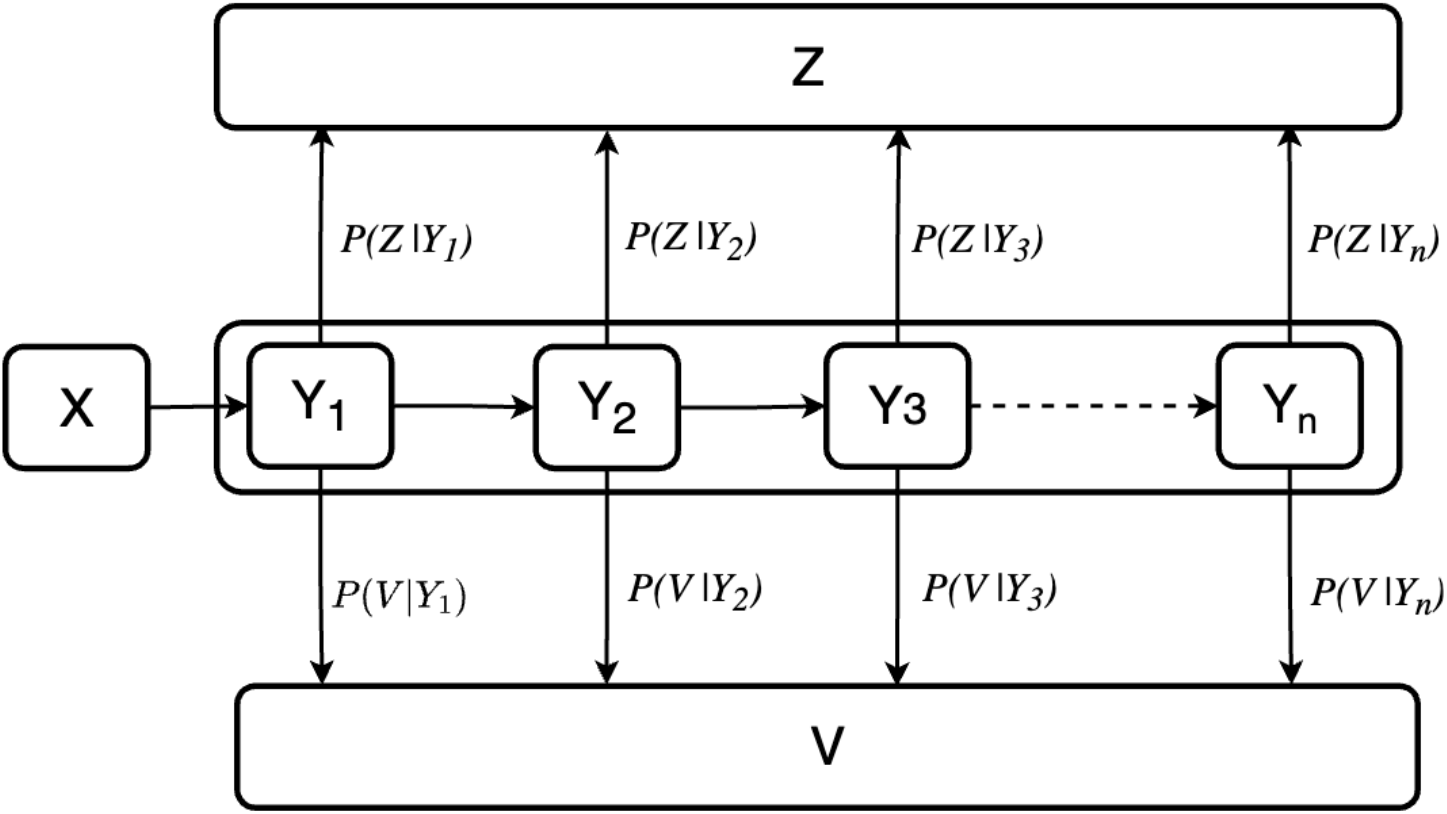
Visualization of the sub-compartment chain in competing risks scenario.

A more detailed discussion regarding competing risks and multinomial, along with their corresponding delay-integro differential equation formulation, is provided in Supplementary Materials 1.

Implementation details and comparison to alternative packages is presented in Supplementary Materials 2.

## 3. Example application

The users can install the package from CRAN by running the following command in R.

`install.packages(“denim”)`

Then load it into R with

`library (denim)`

To create a simulation in denim, the users need to specify (i) model structure (compartments and the transitions between compartments), (ii) initial state of each compartment, (iii) simulation duration and duration for each time step.

In the following sub sections, we will go through a process of building a compartmental model in denim and introduce the core functionalities of the package. A visual presentation for the model being built is shown in Fig. 6. The example was run using denim version 1.2.2.

**Figure 6.**
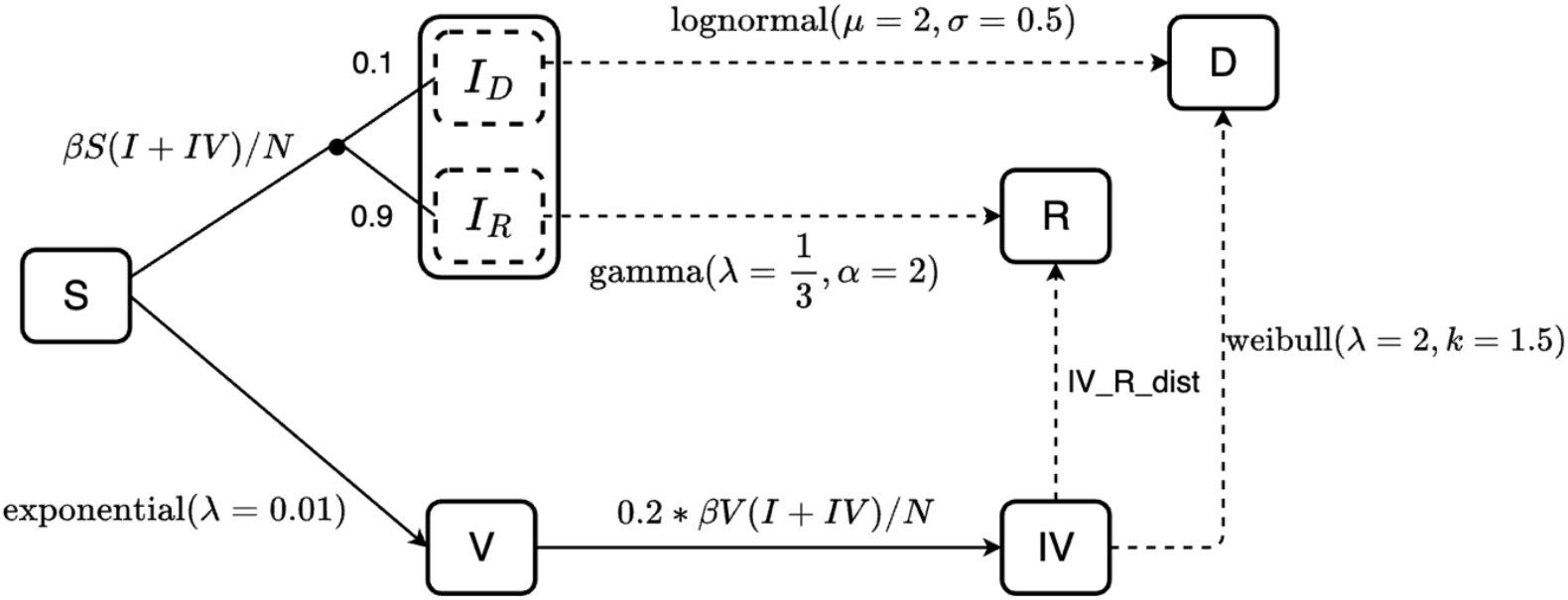
An example of a compartmental model. The dashed lines indicate transitions described by dwell time distributions, while solid lines indicate transition rates. Susceptibles (S) can get infected (I) or vaccinated (V) and these events are treated as competing risks. Among the infected, 90% recover (R) and 10% die (D), with recovery times following a gamma distribution and death times a lognormal distribution. Vaccinated individuals can still be infected (IV). Individuals in IV can either recover or die, following a nonparametric distribution (denoted IV_R_dist in the diagram) and weibull distribution respectively.

**Figure 7.**
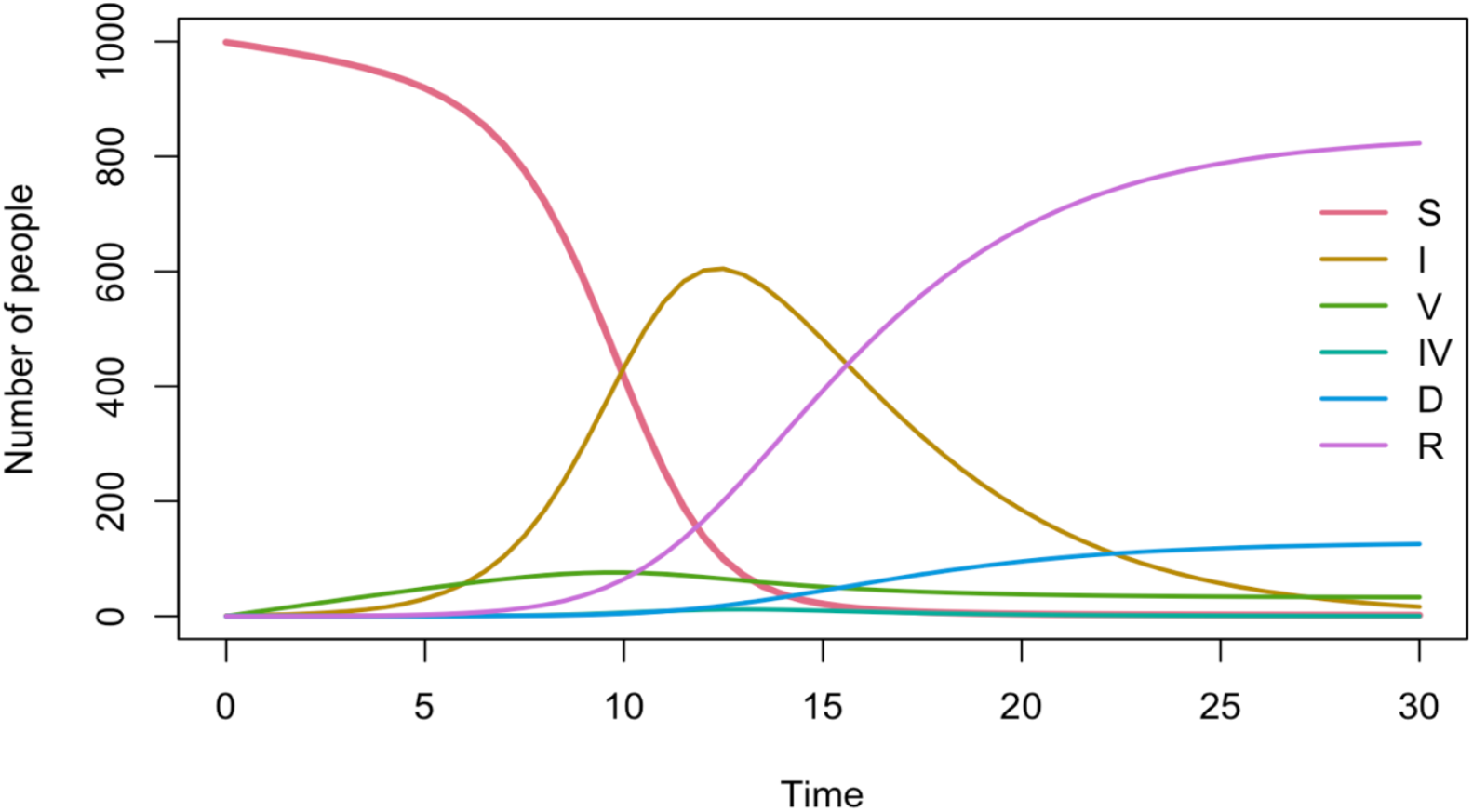
Denim output for the example model.

### 3.1. Model structure

A compartmental model consists of compartments and transitions between them. In denim, model structure is defined by a set of *key*-*value* pairs where: *key* specify the direction of population flow between compartments and *value* is a built-in dwell-time distribution function or mathematical expression to describe that transition.

Table 1 lists all the transitions implemented in denim and their required parameters.

**Table 1:** Built-in transition functions.

| Transition type | Function name | Parameters | Description |
| --- | --- | --- | --- |
| Distribution | d_exponential | rate | for exponentially distributed dwell time |
|  | d_gamma | rate, shape | for gamma distributed dwell time |
|  | d_lognormal | mu, sigma | for lognormal distributed dwell time |
|  | d_weibull | scale, shape | for weibull distributed dwell time |
|  | nonparametric | vector of probabilities or numbers | for user defined dwell time distribution |
| Math expression |  | string for math expressions | for user defined math expression for transition per timestep |

These *key*-*value* pairs can be defined in 2 ways: (i) by using denim domain-specific language (DSL) or (ii) defined as a list in R. This paper will focus on demonstrating the use of denim DSL.

All transitions must be defined in the format of `from -> to = transition`, and code written in denim DSL must be parsed by the function `denim_dsl.` The code for defining the example model is as follows:

~~~
modelStructure <-denim_dsl({
    S -> I = beta * S * (I + IV) / N
    S -> V = d_exponential(0.01)
    0.1 * I -> D = d_lognormal(2, 0.5)
    0.9 * I -> R = d_gamma(1/3, 2)
    V -> IV = 0.2 * beta * V * (I + IV) / N
    IV -> R = nonparametric(iv_r_dist)
    IV -> D = d_weibull(scale = 2, shape = 1.5)
})
~~~

The option to explicitly declare dwell time distribution using built-in functions offers 2 key advantages: (i) from a modeling perspective, denim abstracts away the coding process to implement the desired distribution, allowing modelers more time to focus on the model structure instead; (ii) from a code reviewing perspective, the code’s readability and maintainability is greatly enhanced since transitions are explicitly defined.

Model structure in denim can easily be scaled up by adding more transitions, each of which typically requires only a single line of code. Moreover, model structure can include additional arbitrary parameters as variables for mathematical expressions or distributional parameters (`beta`, `N` and `iv_r_dist` in the example code) which allows customizable transitions.

### 3.2. Model configurations

Initial values for each compartment can be defined as a *key*-*value* pairs (as R named list or named vector) where *key* is the compartment name and *value* is the initial population for the compartment.

Any additional parameters (i.e., variables in math expression transition and distributional parameters) must also be provided in a similar *key*-*value* pairs where *key* is the name of the parameter and *value* is the corresponding value.

Initial state and parameters for the model above can be defined as followed

~~~
initialValues <-c(S = 999, I = 1, R = 0, V = 0, IV = 0, D = 0)
parameters <-list(
  beta = 0.9,
  N = 1000,
  iv_r_dist = c(0, 0.15, 0.15, 0.05, 0.2, 0.2, 0.25)
)
~~~

In situations where there are multiple sub-compartments, by default, the initial population of a compartment is assigned to the first sub-compartment. In this example, denim will internally initialize the first sub-compartment of I→D chain to 0.9, that of I→V chain to 0.1 and the subsequent sub-compartments with population of 0. The user can also choose to distribute initial values based on the waiting time distribution by setting the parameter `dist_init` of distribution transitions to `TRUE` as followed.

~~~
distributeInitVal <-denim_dsl({
    S -> I = beta * S * (I + IV) / N
    S -> V = d_exponential(0.01)
    0.1 * I -> D = d_lognormal(2, 0.5, dist_init = TRUE)
    0.9 * I -> R = d_gamma(1/3, 2, dist_init = TRUE)
    V -> IV = 0.2 * beta * V * (I + IV) / N
    IV -> R = nonparametric(iv_r_dist)
    IV -> D = d_weibull(scale = 2, shape = 1.5)
})
~~~

In competing risks scenario (i.e., `IV -> R` and `IV -> D`), `dist_init` configuration is ignored and the default behavior is applied.

### 3.3. Simulator

After defining model structure and configuration, a simulation can be created by using function **sim** of denim. Aside from the model structure and configurations above, the users need to provide 2 additional parameters `simDuration` for simulation duration and `timeStep` for the duration of each time step.

To simulate the model defined above with duration of 30 and duration for each time step is 0.5, the code in R is as followed

~~~
simulation <-sim(transitions = modelStructure,
           initialValues = initialValues, parameters = parameters,
           simulationDuration = 30,
           timeStep = 0.5)
~~~

### 3.4. Output

By default, denim returns an R 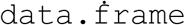 containing population for each compartment per time step. Additionally, the package also provides a built-in visualization tool to quickly plot compartments’ population over time.

Simply call `print(simulation)` to print the 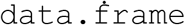, or call `plot(simulation)` to create a plot for the disease dynamics over time.

The output for `plot(simulation, ylim = c(1,1000))`is as followed.

## 4. Discussion

In this paper, we have proposed a modeling framework to easily and intuitively render any sort of dwell time distributions. Accompanying it is the denim package which implements the proposed algorithm, with a user-friendly interface to define model structure in a clear, concise way. We also demonstrated how a compartmental model can be seamlessly translated into code in denim.

We envision that denim will be an essential tool for model selection, thanks to the ease of modification from one dwell time distribution to another. Denim is especially useful for modeling diseases with infectious period distributions that cannot be formulated using ODEs (e.g., log-normal (NISHIURA & EICHNER, 2006), Weibull (Hellewell et al., 2020; Kuk & Ma, 2005), gamma with non-integer shape parameter). Additionally, the declarative nature of the package also enhances the readability of the model definition, thus promoting reproducibility, and enhance the maintainability and reusability of the code.

Compared to other existing packages to handle compartmental model (e.g., deSolve (Soetaert et al., 2008), diffeqr (Rackauckas, 2018)), denim is more computational expensive both in terms of run time and memory space (due to the sub-compartmental structure and iterative nature of the algorithm). Another potential issue to be considered is the discrete time nature of denim, implying that the value of timestep may have an impact on the output of the model. However, for most practical applications, the longer run time remains manageable. These downsides are also compensated by several key advantages:

- Concise model definition: using denim DSL, transition between 2 compartments can be defined in a single line of code. This simplicity is particularly apparent when comparing the implementation for Erlang distributed transition using deSolve by applying Linear Chain Trick (refer to Supplementary Material 3).
- Flexible dwell time distribution: the users can easily incorporate any transition distribution shapes, which is essential for assessing the impact of different dwell time distribution on the disease dynamics. Notably, denim is the only package that can handle multiple competing out-going transitions that are arbitrarily distributed.
- Direct integration of empirical data: the option to provide distribution as a histogram allows the users to directly use empirical data on dwell-time for modeling.

While there exist other simulation tools, to the best of our knowledge, there is no simulation package tailored for simulating compartmental models with diverse built-in dwell-time distributions. To promote adoption, we also provide step-by-step migration guide from deSolve – a widely used ODE based modeling package, to denim (refer to Supplementary Material 3).

For future versions of the package, we plan to provide support for structured compartments (for age, spatial or social structured data) and inclusion of stochasticity to account for uncertainties in disease dynamics. We also plan to extend support for a broader range of parametric distributions, including but not limited to: Pareto, inverse

Gaussian, Gompertz, and log-logistic. Finally, improving the package’s runtime is also one of the priorities, since this can be essential during the model fitting process.

## Acknowledgement

We thank Tuyen Huynh for setting up unit testing for C++ code and Duong Thuy Trang for brainstorming the initial algorithm with us.

Thinh Ong, Anh Phan and Marc Choisy are supported by the Wellcome Trust through the core grant of OUCRU (225167/A/22/Z).

## Author contribution

Marc Choisy, Thinh Ong and Lam Ha conceived the ideas and designed the methodology. Thinh Ong and Anh Phan developed the package. Marc Choisy, Thinh Ong and Anh Phan led the writing of the manuscript. All authors contributed critically to the drafts and gave final approval for publication.

