## Supplementary Material 1 for "denim: an R package for deterministic compartmental models with flexible dwell time distributions"

### Supplementary 1

#### 1 Compartmental model in terms of hazard rate

denim's proposed algorithm was built on the idea of re-formulating the compartmental model in terms of hazard rates. This approach has been explored in a previous work by Tallis (1994).

We first revisit the components of survival analysis under the context of compartmental model.

- $\tau$ is the dwell time, i.e., the time since entering a compartment (also referred to as the secondary time scale in survival analysis).
- $f(\tau)$ is the probability density function (PDF) of the distribution of the dwell time in the compartment and indicates the density of probability that an individual leaves the compartment at time $\tau$.
- $S(\tau)$ is the survival function and indicates the probability that an individual is still in the compartment at time $\tau$. It is worth pointing out that $f(\tau) = -\frac{d}{d\tau}S(\tau)$ or, equivalently, that the survival function is the complement of the cumulative distribution function (CDF): $S(\tau) = 1 - CDF(\tau)$.
- $h(\tau)$ is the hazard rate indicating the instantaneous probability of leaving a compartment, given that the individual is still in the compartment at time $\tau$: $h(\tau) = \frac{f(\tau)}{S(\tau)}$

Functions subscripts denote the corresponding transition. For example, $f_{I \to R}(\tau)$ refers to the PDF for the $I \to R$ transition.

For example, the system of differential equations for an SIR model, written in terms of hazard functions and the dwell time $\tau$, is as followed.

$$\begin{cases} \frac{dS(t)}{dt} = -\int_0^{\mathcal{T}_S} h_{S\to I}(\tau)S(t,\tau)d\tau \\ \frac{dI(t)}{dt} = \int_0^{\mathcal{T}_S} h_{S\to I}(\tau)S(t,\tau)d\tau - \int_0^{\mathcal{T}_I} h_{I\to R}(\tau)I(t,\tau)d\tau \\ \frac{dR(t)}{dt} = \int_0^{\mathcal{T}_I} h_{I\to R}(\tau)I(t,\tau)d\tau \end{cases} \quad (1)$$

Where:

- $\mathcal{T}_S$ and $\mathcal{T}_I$ denotes the maximal time spent in the compartment $S$ and $I$ respectively.
- $S(t,\tau)$ is the sub-population of $S$ at time $t$ that have been in S for a duration $\tau$.

- Similarly, $I(t,\tau)$ is the sub-population of $I$ at time $t$ that have been in I for a duration $\tau$.

It is essential to distinguish $t$ and $\tau$. Here, $t$ refers to the **primary time scale** (the calendar time), while the **secondary time scale** $\tau$ refers to how long individuals have stayed in a compartment at the time $t$.

##### 1.1 Interpretation of traditional SIR model in terms of hazard rate

The traditional formulation for SIR model is typically written as

$$\begin{cases} \frac{dS(t)}{dt} = -\beta \frac{I(t)}{N} S(t) \\ \frac{dI(t)}{dt} = -\beta \frac{I(t)}{N} S(t) - \gamma I(t) \\ \frac{dR(t)}{dt} = \gamma I(t) \end{cases}$$

Drawing connection between $h_{I\to R}(\tau)$ and $\gamma$ is relatively straight-forward when we consider the assumption that the duration of the infectious period is exponentially distributed with rate $\gamma$.

$$h_{I\to R}(\tau) = \frac{f_{I\to R}(\tau)}{S_{I\to R}(\tau)} = \frac{\gamma e^{-\gamma\tau}}{e^{-\gamma\tau}} = \gamma$$

$\frac{dR(t)}{dt}$ in Equation 1 can then be rewritten as followed

$$\frac{dR(t)}{dt} = \int_0^{\mathcal{T}_I} \gamma\, I(t,\tau) d\tau = \gamma \int_0^{\mathcal{T}_I} I\,(t,\tau) d\tau = \gamma I(t)$$

Similarly, we can draw the connection that $h_{S\to I}(\tau) = \beta \frac{I(t)}{N}$. Since $h_{S\to I}(\tau)$ only varies with respect to $t$ and not $\tau$, formulation for $\frac{dS(t)}{dt}$ in Equation 1 can then be simplified as followed.

$$\frac{dS(t)}{dt} = -\int_0^{\mathcal{T}_S} \beta \frac{I(t)}{N} S(t,\tau) d\tau = -\beta \frac{I(t)}{N} \int_0^{\mathcal{T}_S} S\,(t,\tau) d\tau = -\beta \frac{I(t)}{N} S(t)$$

This demonstrates that the force of infection $\beta \frac{I(t)}{N}$ and recovery rate $\gamma$ can be interpreted as the hazard rate for $S \to I$ event and $I \to R$ event respectively.

##### 1.2 Drawing connection to denim's algorithm

Recall the problem set up from the main paper, where we consider a simple $X \to Y \to Z$ model, with emphasis on the transition $Y \to Z$. We can interpret the proposed algorithm as a discrete approximation of the following system of integro-differential equations.

$$\begin{cases} \dfrac{dX(t)}{dt} = -N_Y(t) \\ \dfrac{dY(t)}{dt} = N_Y(t) - \displaystyle\int_0^{\mathcal{T}_Y} h_{Y\to Z}(\tau) Y(t,\tau) d\tau \\ \dfrac{dZ(t)}{dt} = \displaystyle\int_0^{\mathcal{T}_Y} h_{Y\to Z}(\tau) Y(t,\tau) d\tau \end{cases} \quad (2)$$

Where:

- $N_Y(t)$ is the population of $X$ that transition to $Y$ at time $t$.
- $Y(t,\tau)$ is the sub-population of $Y$ at time $t$ that have been in $Y$ for a duration $\tau$ (i.e., $Y(t,\tau) = S_{Y\to Z}(\tau) N_Y(t - \tau)$).

Visualization of the algorithm to handle arbitrary distributed dwell time for $Y$ is presented below.

With: $q_i = \int_{(i-1)\Delta t}^{i\Delta t} h_{Y\to X}(\tau) d\tau = \frac{\int_{(i-1)\Delta t}^{i\Delta t} f_{Y\to X}(\tau) d\tau}{1 - \int_0^{(i-1)\Delta t} f_{Y\to X}(\tau) d\tau} \approx \frac{p_i}{1 - \sum_{j=1}^{i-1} p_i}$

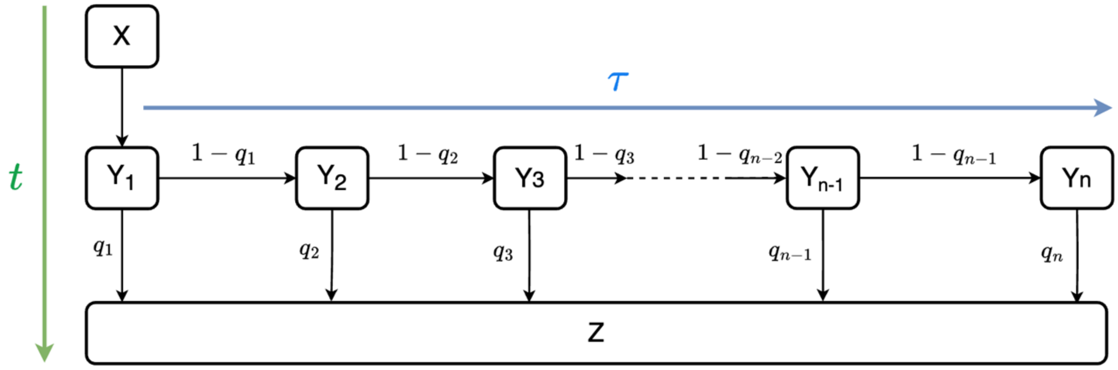


*Figure 1: sub-compartment structure in denim*

Components of this diagram can be interpreted in terms of Equation 2 as followed

- The dwell time distribution of the $Y$ compartment $P_Y = p_1, p_2, \dots, p_n$ where $\sum_{i=1}^{n} p_i = 1$ and $n$ is the maximal dwell time (in time steps) is a discrete approximation for $f_{Y\to Z}(\tau)$: $\mathrm{p_i} = \int_{(\mathrm{i}-1)\Delta\mathrm{t}}^{\mathrm{i}\Delta\mathrm{t}} \mathrm{h_{Y\to Z}}(\tau)\mathrm{d}\,\mathrm{p_i} = \int_{(\mathrm{i}-1)\Delta\mathrm{t}}^{\mathrm{i}\Delta\mathrm{t}} \mathrm{h_{Y\to Z}}(\tau)\mathrm{d}\,\tau$ where $\Delta t$ is the time step.
- $1 - \sum_{j=1}^{i-1} p_j$ representing the probability of staying in $Y$ for at least $i$ time steps is an approximation for $S_{Y\to Z}(\tau)$.
- The transition proportion $q_i = P(Z|Y_i) = \frac{p_i}{1-\sum_{j=1}^{i-1} p_j}$ thus approximates the hazard rate $h_{Y\to Z}(\tau) = \frac{f_{Y\to Z}(\tau)}{S_{Y\to Z}(\tau)}$.

- Sub-compartments $Y_i$ corresponds to individuals that have stayed in $Y$ for $i$ time steps, effectively representing the discrete version of $Y(t, \tau)$ where $\mathcal{T}_Y$[1] is discretized by time step.

- Thus, the total outgoing population for $Y$ computed by $\sum_{i=1}^{n} q_i Y_i$ is a discrete approximation for $\int_0^{\mathcal{T}_Y} h_{Y \to Z}(\tau) Y(t, \tau) d\tau$.

#### 2 Formulating multiple outgoing compartments

There are 2 main approaches offered by denim to formulate multiple outgoing compartments:

- As competing risks

- Or as multinomial transitions

The difference between the two comes from the type of data / information we have at hand. Consider a scenario where compartment $X$ can transition to $n$ out compartments denoted $O^{(i)}$ where $i \in \{1,2, \dots, n\}$. The data we can get include:

1. The distribution of dwelling time in compartment $X$, denoted $f_X(\tau)$
2. The distributions of dwelling time in compartment X respective to each of the $O^{(i)}$ compartments that are ultimately reached upon exiting X, denoted $f_{X \to O^{(i)}}(\tau)$
3. The proportion of individuals in compartment $X$ that ends up in each $O^{(i)}$, denoted $w_i$ where $\sum_{i=1}^{n} w_i = 1$

Transitions are modeled as competing risks if we have information 2, and modeled as multinomial transitions if we have information 2+3 or 1+3. A diagram illustrates these scenarios when $n = 2$ is provided below.

---

[1] The most intuitive choice for $\mathcal{T}_Y$ is the value where $F_{Y \to Z}(\mathcal{T}_Y) = 1$ with $F_{Y \to Z}(\tau)$ being the cumulative distribution (CDF) for $Y \to Z$ transition. However, for most parametric distributions, the cumulative distribution is asymptotic to 1 thus $\mathcal{T}_Y$ is usually set to $t$ hence the upper limit for $\mathcal{T}_Y$ is $\max(t)$ i.e., calendar time. For denim in particular, $\mathcal{T}_Y$ is set to a value where the CDF is sufficiently close to 1, i.e., $F_{Y \to Z}(\mathcal{T}_Y) \geq 1 -$ error tolerance. Under this approach, in most practical use cases we would get $\max(t) > \mathcal{T}_Y$ which makes it a more efficient way to generate the sub-compartment chain.

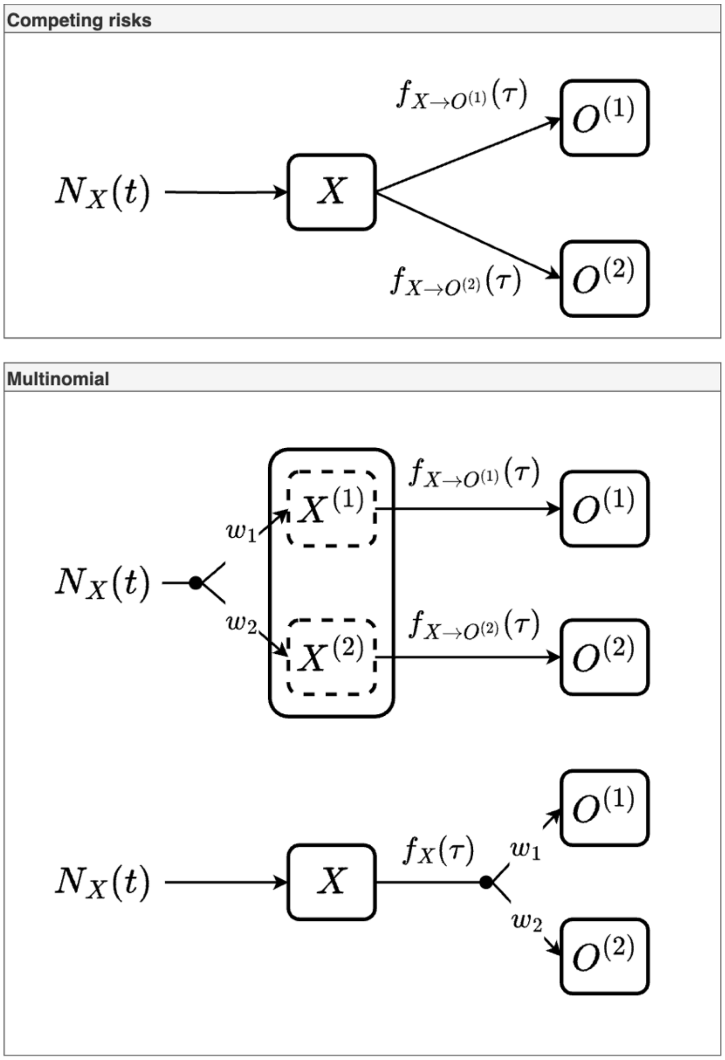


*Figure 2: 3 scenarios for modeling multiple outgoing compartments, depending on the information available. The lines extending from the black dot indicate the proportion of individuals that eventually enter each outgoing compartment at equilibrium.*

#### 2.1 Competing risks

Under this scenario, we model the out-going transitions as $n$ independent, competing events each with its own arbitrary distribution. The definition of competing risks here closely follows that under survival analysis context, which entails the following:

- Individuals are susceptible to all $n$ events, however, one can only experience 1 event. For example, one individual can either transition to $O^{(1)}$ or $O^{(2)}$ but not both.
- By extension, individual will only experience the first event that occur. For example, let $T_{X\to O^{(1)}}$ and $T_{X\to O^{(2)}}$ be 2 random variables for the times at which an individual transition to $O^{(1)}$ and $O^{(2)}$ respectively. The time at which the individual leaves $X$ is then given by: min $(T_{X\to O^{(1)}}, T_{X\to O^{(2)}})$.

Formulating this model in terms of hazard rate is straightforward when we consider the fact that when competing risks are independent, the overall hazard rate is simply the sum of all the hazard rates for each risk, represented as the following equation (refer to Section 3.1 for proof).

$$h_X(\tau) = \sum_{i=1}^{n} h_{X\to O^{(i)}}(\tau)$$

Where $h_X(\tau)$ is the overall hazard rate of leaving $X$ at time $\tau$.

The formulation for the proposed scenario can then be represented by:

$$\begin{cases} \dfrac{dX(t)}{dt} = N_X(t) - \displaystyle\int_0^{\mathcal{T}_X} h_X(\tau)X(t,\tau)d\tau = N_X(t) - \int_0^{\mathcal{T}_X} \sum_{i=1}^{n} h_{X\to O^{(i)}}(\tau)X(t,\tau)d\tau \\ \dfrac{dO^{(1)}(t)}{dt} = \displaystyle\int_0^{\mathcal{T}_X} h_{X\to O^{(1)}}(\tau)\, X(t,\tau)d\tau \\ \dots \\ \dfrac{dO^{(n)}(t)}{dt} = \displaystyle\int_0^{\mathcal{T}_X} h_{X\to O^{(n)}}(\tau)\, X(t,\tau)d\tau \end{cases}$$

Where $N_X(t)$ is the in-coming population of $X$ at time $t$.

Note that $h_{X\to O^{(i)}}(\tau)X(t,\tau) = h_{X\to O^{(i)}}(\tau)S_X(\tau)N_X(t-\tau)$ and $h_{X\to O^{(i)}}(\tau)S_X(\tau)$ is what referred to as Cumulative Incidence Function (CIF) in survival analysis.

A diagram to demonstrate the discrete representation for the formula when $n = 2$ is provided below

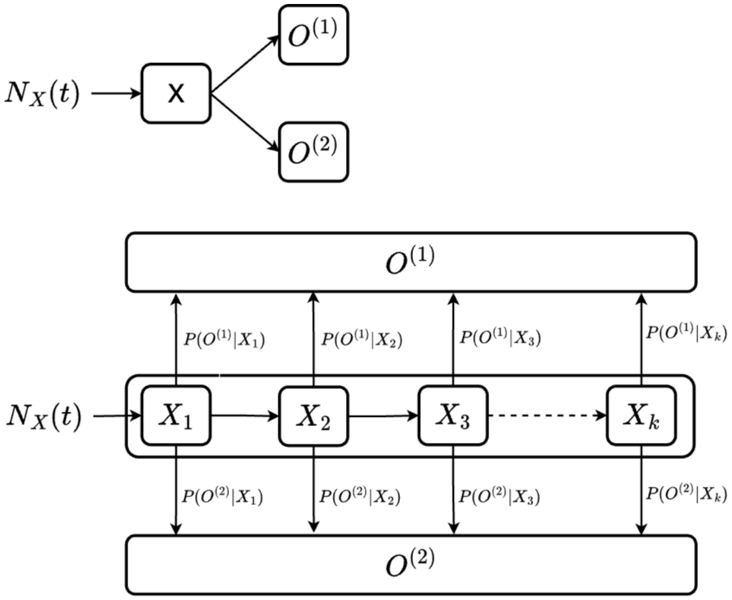


*Figure 3: Discrete representation for competing risks. Upper figure demonstrates the overall flow between compartments, while the lower figure expands on this by showing the sub-*

*compartments structure within $X$ compartment. $P\left(O^{(1)}|X_i\right)$ and $P\left(O^{(2)}|X_i\right)$ are the discrete estimation for $h_{X\to O^{(1)}}(\tau)$ and $h_{X\to O^{(2)}}(\tau)$ respectively.*

#### 2.2 Multinomial

Consider a scenario where compartment $X$ can transition to $n$ out compartments denoted $O^{(i)}$ where $i \in \{1,2,\dots,n\}$. We want to model such that the population distributed across $\{O^{(1)}, O^{(2)}, \dots, O^{(n)}\}$ compartments following the distribution $\{w_1, w_2, \dots, w_n\}$ where $\sum_{i=1}^{n} w_i = 1$.

There are 2 main approaches to achieve this

**Approach 1:** Upon *exiting* $X$, the out-going population is distributed across $n$ out-compartments based on the specified distribution. Under this approach, individuals in $X$ share the same dwell time distribution.

This approach can be represented using the following system of derivatives.

$$\begin{cases} \dfrac{dX(t)}{dt} = N_X(t) - \displaystyle\int_0^{\mathcal{T}_X} h_X(\tau)X(t,\tau)d\tau \\ \dfrac{dO^{(1)}(t)}{dt} = w_1 \displaystyle\int_0^{\mathcal{T}_X} h_X(\tau)X(t,\tau)d\tau \\ \dots \\ \dfrac{dO^{(n)}(t)}{dt} = w_n \displaystyle\int_0^{\mathcal{T}_X} h_X(\tau)X(t,\tau)d\tau \end{cases}$$

Where $N_X(t)$ is the in-coming population of $X$ at time $t$.

A diagram to demonstrate the discrete representation for the formula when $n = 2$ is provided below

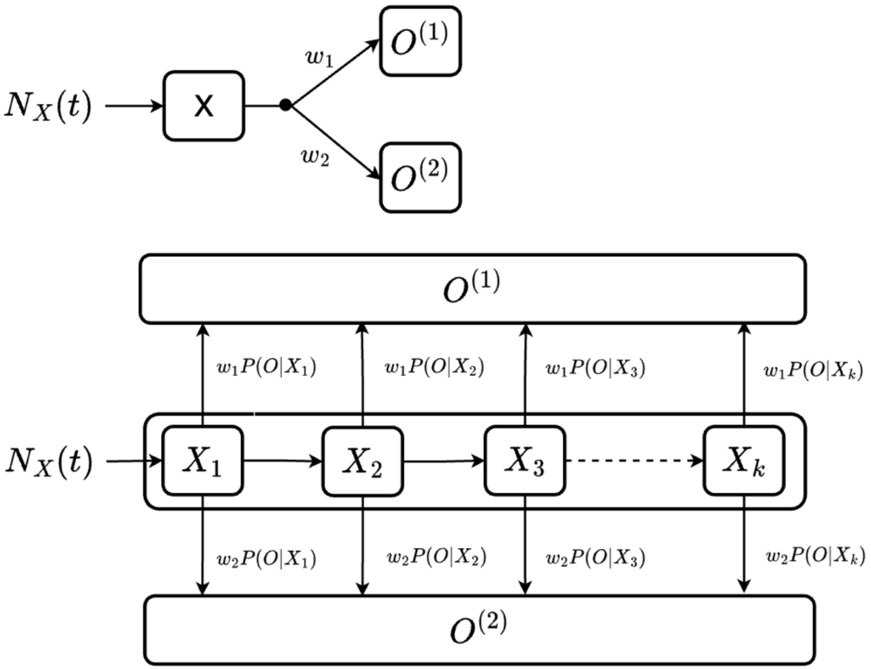


*Figure 4: Discrete representation for Approach 1. Upper figure demonstrates the overall flow between compartments, while the lower figure expands on this by showing the sub-compartments structure within $X$ compartment. $P(O|X_i)$ is the discrete estimation for $h_X(\tau)$.*

**Approach 2:** Upon *entering* $X$, in-coming population is distributed across $n$ different processes to reach $O^{(i)}$, following the specified distribution. Under this approach, individuals in $X$ are partitioned to follow $n$ dwell time distributions depending on which $O^{(i)}$ they are transitioning to. This approach can be represented as followed

$$\begin{cases} \dfrac{dX(t)}{dt} = N_X(t) - \displaystyle\sum_{i=1}^{n} \int_0^{\mathcal{T}_{X\to O^{(i)}}} h_{X\to O^{(i)}}(\tau)\, X^{(i)}(t,\tau) d\tau \\ \dfrac{dO^{(1)}(t)}{dt} = \displaystyle\int_0^{\mathcal{T}_{X\to O^{(1)}}} h_{X\to O^{(1)}}(\tau)\, X^{(1)}(t,\tau) d\tau \\ \dots \\ \dfrac{dO^{(n)}(t)}{dt} = \displaystyle\int_0^{\mathcal{T}_{X\to O^{(n)}}} h_{X\to O^{(n)}}(\tau) X^{(n)}(t,\tau) d\tau \end{cases} \tag{3}$$

Where:

- $N_X(t)$ is the in-coming population of $X$ at time $t$.
- $X^{(i)}$ denotes the sub-population of $X$ that is transitioning to $O^{(i)}$.
- $\mathcal{T}_{X\to O^{(i)}}$ is the maximal time to transition from $X$ to $O^{(i)}$.

Note that $X^{(i)}$ can be rewritten in terms of $N_X$ as followed

$$X^{(i)}(t,\tau) = X^{(i)}(t-\tau,0)\, S_{X\to O^{(i)}}(\tau) = w_i N_X(t-\tau) S_{X\to O^{(i)}}(\tau)$$

[Equation 3](#) can thus be rewritten as

$$
\begin{cases}
\dfrac{dX(t)}{dt} = N_X(t) - \displaystyle\sum_{i=1}^{n} \int_0^{\mathcal{T}_{X\to O^{(i)}}} w_i \; N_X(t-\tau)\, f_{X\to O^{(i)}}(\tau) d\tau \\
\dfrac{dO^{(1)}(t)}{dt} = \displaystyle\int_0^{\mathcal{T}_{X\to O^{(1)}}} w_1 \, N_X(t-\tau)\, f_{X\to O^{(1)}}(\tau) d\tau \\
\dots \\
\dfrac{dO^{(n)}(t)}{dt} = \displaystyle\int_0^{\mathcal{T}_{X\to O^{(n)}}} w_n \; N_X(t-\tau)\, f_{X\to O^{(n)}}(\tau) d\tau
\end{cases}
\tag{4}
$$

A diagram to illustrate the discrete representation for Equation 3 when $n = 2$ is provided below

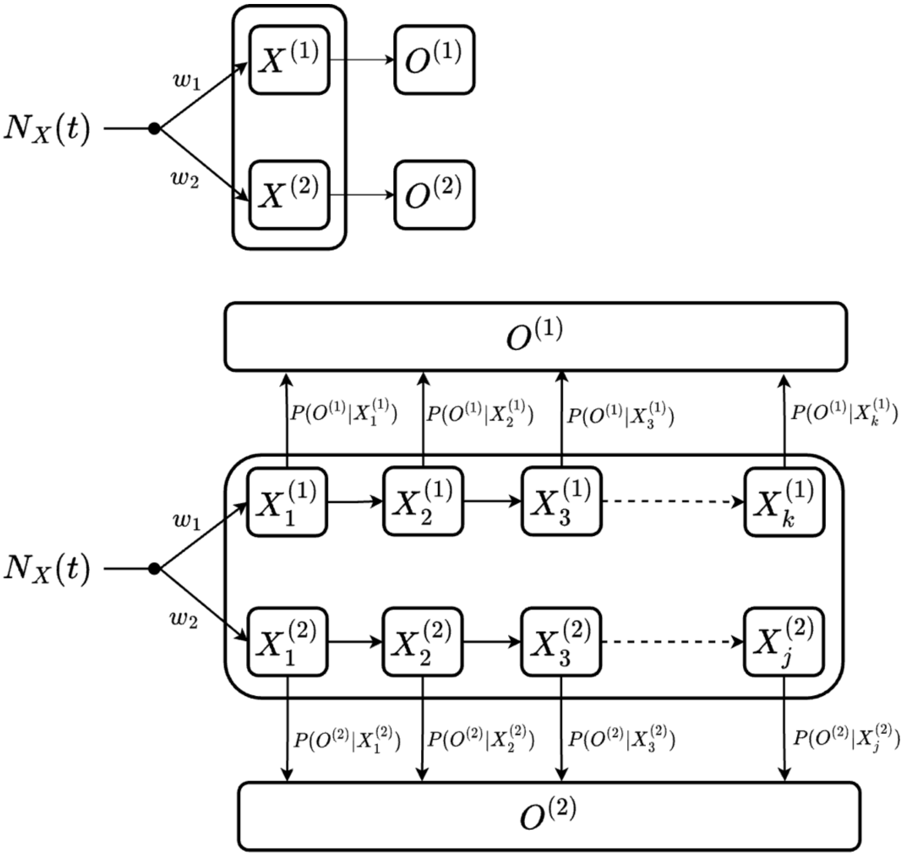


*Figure 5: Discrete representation of Approach 2. Upper figure demonstrates the overall flow between compartments, note that compartment $X$ is partitioned into $X^{(1)}$ and $X^{(2)}$ which will transition to $O^{(1)}$ and $O^{(2)}$ respectively. The lower figure expands on this by showing sub-compartments structures of $X^{(1)}$ and $X^{(2)}$. $P\left(O^{(1)}|X_i^{(1)}\right)$ and $P\left(O^{(2)}|X_i^{(2)}\right)$ are discrete estimations for $h_{X\to O^{(1)}}(\tau)$ and $h_{X\to O^{(2)}}(\tau)$ respectively.*

Section 3.2 below demonstrates that Approach 1 is equivalent to a special case of Approach 2, where the dwell time distributions $f_{X\to O^{(1)}}(\tau), f_{X\to O^{(2)}}(\tau), \dots, f_{X\to O^{(n)}}(\tau)$ are identical. Thus, Approach 2 is how multinomial is implemented in denim.

### 3 Appendix

#### 3.1 Proof 1

Recall that $S_X(\tau)$ is equivalent to $P(T_X > \tau)$ (i.e., probability that individuals stay in $X$ compartment for at least $\tau$ time period).

$$P(T_X > \tau) = P\left(T_{X\to O^{(1)}} > \tau, T_{X\to O^{(2)}} > \tau, \dots, T_{X\to O^{(n-1)}} > \tau, T_{X\to O^{(n)}} > \tau\right)$$

Under the assumption that these competing risks are independent

$$P(T_X > \tau) = P\left(T_{X\to O^{(1)}} > \tau\right)P\left(T_{X\to O^{(2)}} > \tau\right)\dots P\left(T_{X\to O^{(n-1)}} > \tau\right)P\left(T_{X\to O^{(n)}} > \tau\right)$$

$$S_X(\tau) = P(T_X > \tau) = \prod_{i=1}^{n} P\left(T_{X\to O^{(i)}} > \tau\right)$$

The hazard function for leaving $X$ can then be computed by

$$h_X(t) = \frac{-\frac{d}{dt}\prod_{i=1}^{n} S_{X\to O^{(i)}}(t)}{\prod_{i=1}^{n} S_{X\to O^{(i)}}(t)}$$

By applying the product rule for differentiation, we have

$$h_X(t) = \frac{-\sum_{i=1}^{n}\frac{d}{dt}S_{X\to O^{(i)}}(t)\ \prod_{j=1}^{n,n\neq i} S_{X\to O^{(j)}}(t)}{\prod_{i=1}^{n} S_{X\to O^{(i)}}(t)}$$

Where $\prod_{j=1}^{n,n\neq i} S_{X\to O^{(j)}}(t)$ denotes product of $S_{X\to O^{(j)}}(t)$ where $j$ ranges from 1 to $n$ and $j \neq i$. (i.e., $j \in \{1,2,3,\dots,n\}\setminus\{i\}$)

Recall that $f(t) = -\frac{d}{dt}S(t)$

We can then rewrite $h_X(t)$ as followed

$$h_X(t) = -\sum_{i=1}^{n}\frac{-f_{X\to O^{(i)}}(t)\ \prod_{j=1}^{n,n\neq i} S_{X\to O^{(j)}}(t)}{\prod_{k=1}^{n} S_{X\to O^{(k)}}(t)}$$

Since $\frac{\prod_{j=1}^{n,n\neq i} S_{X\to O^{(j)}}(t)}{\prod_{k=1}^{n} S_{X\to O^{(k)}}(t)} = \frac{\prod_{j=1}^{n,n\neq i} S_{X\to O^{(j)}}(t)}{S_{X\to O^{(i)}}(t)\prod_{k=1}^{n,n\neq i} S_{X\to O^{(k)}}(t)} = \frac{1}{S_{X\to O^{(i)}}(t)}$

We can further simplify $h_X(t)$ as

$$h_X(t) = -\sum_{i=1}^{n} \frac{-f_{X\to O^{(i)}}(t)}{S_{X\to O^{(i)}}(t)} = \sum_{i=1}^{n} \frac{f_{X\to O^{(i)}}(t)}{S_{X\to O^{(i)}}(t)} = \sum_{i=1}^{n} h_{X\to O^{(i)}}(t)$$

##### 3.2 Proof 2

From the system of derivatives for Approach 1

$$\begin{cases} \dfrac{dX(t)}{dt} = N_X(t) - \displaystyle\int_0^{\mathcal{T}_X} h_X(\tau)\, X(t,\tau) d\tau \\ \dfrac{dO^{(1)}(t)}{dt} = w_1 \displaystyle\int_0^{\mathcal{T}_X} h_X(\tau)\, X(t,\tau) d\tau \\ \dots \\ \dfrac{dO^{(n)}(t)}{dt} = w_n \displaystyle\int_0^{\mathcal{T}_X} h_X(\tau)\, X(t,\tau) d\tau \end{cases}$$

We can rewrite $X(t,\tau)$ as $X(t,\tau) = X(t-\tau,0)\, S_X(\tau) = N_X(t-\tau)\, S_X(\tau)$

$N_X(t-\tau)$ can also be expressed as $N_X(t-\tau) = \sum_{i=1}^{n} w_i\, N(t-\tau)$

Substituting these to the system yields

$$\begin{cases} \dfrac{dX(t)}{dt} = N_X(t) - \displaystyle\sum_{i=1}^{n} \int_0^{\mathcal{T}_X} w_i\, N(t-\tau)\, f_X(\tau) d\tau \\ \dfrac{dO^{(1)}(t)}{dt} = \displaystyle\int_0^{\mathcal{T}_X} w_1\, N_X(t-\tau)\, f_X(\tau) d\tau \\ \dots \\ \dfrac{dO^{(n)}(t)}{dt} = \displaystyle\int_0^{\mathcal{T}_X} w_n\, N_X(t-\tau)\, f_X(\tau) d\tau \end{cases}$$

Which is equivalent to Equation 4 under the condition that all $X \to O^{(i)}$ transitions share the same dwell time distribution

$$f_{X\to O^{(1)}}(\tau) = f_{X\to O^{(2)}}(\tau) = \dots = f_{X\to O^{(n)}}(\tau) = f_X(\tau)$$
