## Supplementary Material 2 for "denim: an R package for deterministic compartmental models with flexible dwell time distributions"

### 1. Implementation details

#### 1.1. Main components

Internally, the denim package consists of 4 main components: Transition, Compartment, Model and Simulator.

**Transition** computes the maximum dwell time and transition proportion (i.e., $q_i$ in the main text) based on users' input. The denim package offers 2 ways of defining a transition between compartments: (i) a distribution of dwell time (using a parametric distribution or a histogram) and (ii) the number of people transitioning, defined by a math expression. Maximum dwell time for each type of transition is computed as followed:

- parametric distribution: maximum dwell time is the time point $t$ where cumulative probability distribution ($F(t)$) is sufficiently close to 1 (the threshold is $1 -$ error tolerance where error tolerance is an adjustable parameter).
- nonparametric distribution (histogram of a distribution): maximum dwell time is $s$ time steps where $s$ is the length of the user defined probability distribution (i.e., number of bins of the histogram).
- for transition defined using a math expression, the length of sub-compartment is set to 1, and it will be updated by the specified value (evaluated from the expression) every time step.

**Compartment** component will then use the maximum dwell time previously computed to create sub-compartments. It also handles in-coming compartments, out-going compartments and updates its sub-compartments after each time step.

**Model** component manages all the high-level configurations (initial states, transition distributions and their parameters) and instantiate Compartments and Transitions accordingly. It also handles the order in which compartments are updated.

Finally, **Simulator** is where the simulation process takes place. Its core functionalities are to instantiate the model from user's input and to run the simulation process.

#### 1.2. Implementation architecture

For better computation efficiency, the backend of denim is implemented in C++ (i.e., user defined models are interfaced to C++ for the simulation process).

**Transition**, **Compartment**, **Model** components are implemented as classes in C++, while **Simulator** component is a C++ function that returns a DataFrame object and interfaced to R through Rcpp.

User defined model specifications in R are converted to a JSON format, which is then parsed using JSON library in C++ (Lohmann, 2025). Additionally, muParser C++ library (Ingo, n.d.) is used for handling math expression transitions.

### 2. Comparison to alternative solutions

To the best of our knowledge, no other existing package serves the same purpose as denim: to create scalable compartmental models with diverse built-in dwell time distributions. Therefore, we will compare denim to other existing solutions from two perspectives: as a tool to build compartmental models and as an approach to incorporate arbitrary distribution.

Amongst other modeling tools, deSolve (Soetaert et al., 2008) is one of the most widely used R packages. The package, at its core, is an optimized ODE solver making it a popular tool for modelers following the traditional formulation of compartmental models. This, however, means that using deSolve (or similar solvers, for example, diffeqr (Rackauckas, 2018)) also comes with the limitations of ODEs formulation with the main disadvantage being the difficulty of including distribution of waiting time resulting from a non-Markovian process. Another important aspect about denim is that the model definition is much more concise compared to that using deSolve (Soetaert et al., 2008), especially when Linear Chain Trick (Hurtado & Kirosingh, 2019) is involved.

Another popular package for modeling, odin (FitzJohn, 2019), also shares the same limitations as deSolve when defining models as ODEs. However, odin also provides an option for defining discrete models, which makes it possible to implement denim's proposed algorithm. Nevertheless, this approach would require much more complex code, as the users must manually implement the algorithm. Package odin also offers several additional features including the option to create stochastic models and multi-dimensional compartments (for compartment stratification such as age structure), both of which are features not yet implemented in denim.

In terms of methodology, uSEIR (Hernández et al., 2021) also offers an alternative compartmental model formulation which aims to include general distributed dwell-time. Their implementation also makes use of a sub-compartment structures similar to denim. However, there are several key differences between the approaches of uSEIR and denim.

- Model structure: uSEIR was designed for SEIR model, while denim's generalized framework is applicable to any compartmental model structure.
- Transition time distribution: uSEIR specifically targets parametric distributions (exponential, Poisson, gamma), while denim extends its capability to also handle non-parametric distributions.

IONISE (Hong et al., 2024) proposes a similar formulation using delay-integro differential equations, but implements a custom numerical integrator to solve the system of equations, and support wider range of dwell-time distributions (exponential, gamma, inverse-gamma, lognormal, weibull). However, some downsides similar to uSEIR still persist: (i) the model structured is specifically SEIR and cannot be easily scaled up, and (ii) dwell time distributions are still limited to the package built-in parametric ones.

#### 2.1. Benchmark

To assess denim's performance, we benchmark it against other approaches/packages including deSolve (Soetaert et al., 2008), uSEIR implementation in pure Python and

Cython (Hernández et al., 2021), IONISE (Hong et al., 2024), and diffeqr (Rackauckas, 2018).

Fig. 1 visualizes the model structure for benchmarking. The structure closely follows the formulation for the uSEIR model with gamma distributed infectious and recovery time. For model configuration, simulation duration is set to 180 and the duration of the time step is 0.01.

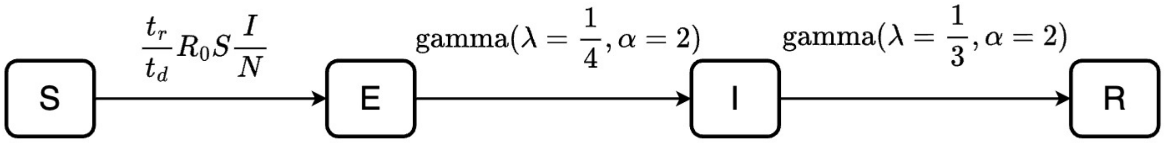


*Figure 1. SEIR model used for benchmarking. Where $R_0$ is the reproduction number, $t_r$ is the expected dwell time in $I$ compartment, $t_d$ is the duration of the time step.*

Table 1 shows the median run time (calculated from 50 runs) of each approach and the variance between runs. The benchmarking process was conducted on a MacBook Pro with M2 Pro chip, 16 GBs of RAM and 10 cores (6 performance and 4 efficiency). The code for benchmarking and additional outputs are available on the denim website (*https://anonymous.4open.science/w/denim-72E0/articles/denim_benchmark.html*) and Github (*https://anonymous.4open.science/r/denim-72E0/vignettes/denim_benchmark.Rmd*) .

*Table 1. Runtime comparison between different approaches.*

| Approach | Backend implementation | Median time (in seconds) | Variance (in seconds) |
|---|---|---|---|
| denim | C++ | 0.7617 | 1.5910e-04 |
| denim (using nonparametric()) | C++ | 1.2232 | 0.0067 |
| odin (discrete-time implementation) | C | 0.3264 | 0.0011 |
| uSEIR (pure python implementation) | Python | 53.7091 | 3.3785 |
| uSEIR (cython implementation) | C | 0.4189 | 8.9641e-05 |
| IONISE | R | 0.0516 | 1.2749e-04 |
| diffeqr (model definition in R) | Julia | 0.0021 | 1.6974e-07 |
| diffeqr (model definition in Julia) | Julia | 1.0252e-4 | 4.7504e-10 |
| deSolve (model definition in R) | Fortran | 0.0034 | 2.8770e-06 |
| deSolve (model definition in C) | C | 1.2671e-4 | 1.8830e-09 |

| Approach | Backend implementation | Median time (in seconds) | Variance (in seconds) |
|---|---|---|---|
| deSolve (model definition in Fortran) | Fortran | 1.7898e-4 | 4.4677e-09 |

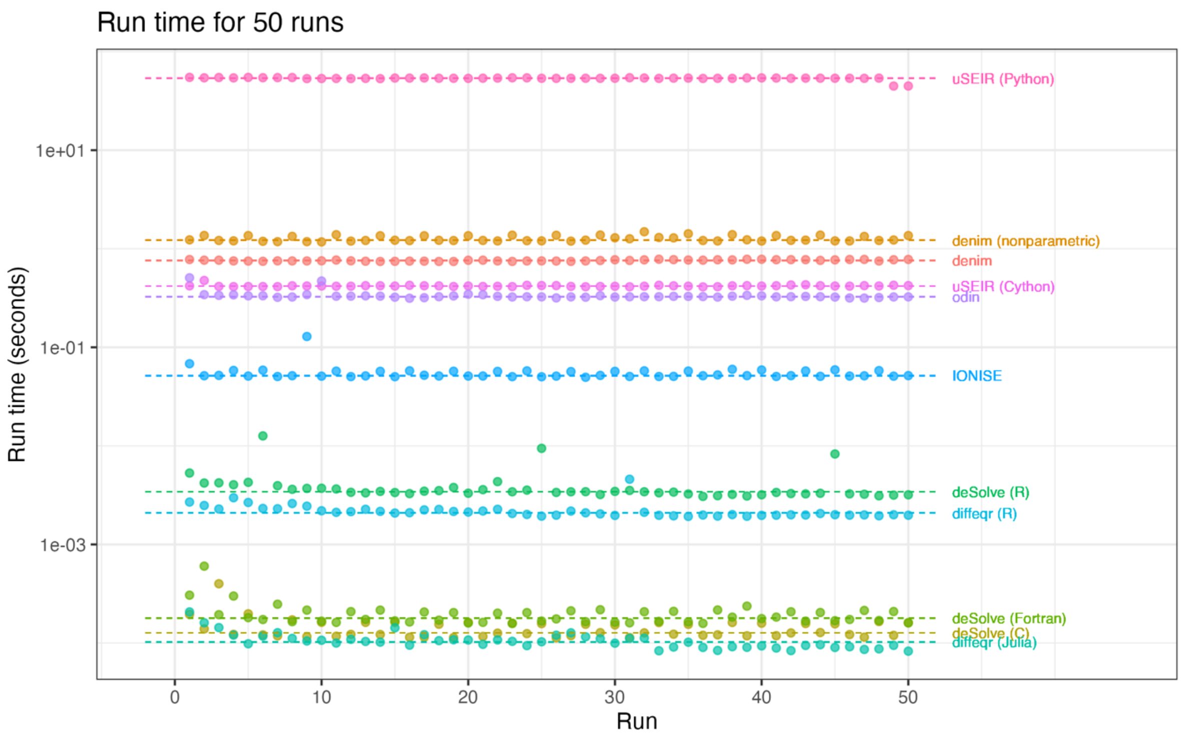


*Figure 2. Visualization of run time for each of the 50 runs. The dashed horizontal line represents the median run time of each approach. Run time is presented in log-scale to better demonstrate difference in run time for faster approaches.*

Unsurprisingly, optimized ODE solvers (deSolve, diffeqr) run significantly faster than alternative approaches, especially when the model is implemented in the solver's native language (Fortran/C for deSolve and Julia for diffeqr). IONISE with its numerical integrator implemented in R cannot achieve comparable speed, however, it comes with the benefit of allowing more flexible dwell time distributions for the transitions.

Among discrete time approaches with similar sub-compartment structure, denim performs slightly worse than uSEIR (with Cython) and odin. This performance gap is likely due to the additional overhead from parsing the model structure during the R–C++ interfacing process. Denim also offers additional advantages of allowing more flexible model structure, and the ease of implementing more complex systems.
