## Supplementary Material 3 for "denim: an R package for deterministic compartmental models with flexible dwell time distributions"

### deSolve to denim

#### Migrate deSolve code to denim

##### Original code in deSolve

The model used for demonstrating the process of migrating code from `deSolve` to `denim` is as followed

```
library(denim)

library(deSolve)

# --- Model definition in deSolve
transition_func <- function(t, state, param){
  with(as.list( c(state, param) ), {
      dS = -beta*S*I/N
      dI1 = beta*S*I/N - rate*I1
      dI2 = rate*I1 - rate*I2
      dI =  dI1 + dI2
      dR = rate*I2
      list(c(dS, dI, dI1, dI2, dR))
  })
}

# ---- Model configuration
parameters <- c(beta = 0.3, rate = 1/3, N = 1000)
initialValues <- c(S = 999, I = 1, I1 = 1, I2=0, R=0)

# ---- Run simulation
times <- seq(0, 100) # simulation duration
ode_mod <- ode(y = initialValues, times = times, parms = parameters, func =
transition_func)
ode_mod <- as.data.frame(ode_mod)
```

##### Model definition

Similar to deSolve, transitions between compartments in denim can be defined using ordinary differential equations (ODEs). However, denim extends it by providing the option to directly express the dwell time distribution.

To utilize this option from denim, the user must first identify which transitions can best describe the ones in their deSolve model.

```
# --- Model definition in deSolve
transition_func <- function(t, state, param){
  with(as.list( c(state, param) ), {
```

```
      # For S -> I transition, since it involves parameters (beta, N),
      # the best transition to describe this is using a mathematical formula
      dS = -beta*S*I/N

      # For I -> R transition, linear chain trick is applied --> implies
Erlang distributed dwell time
      # Hence, we can use d_gamma from denim
      dI1 = beta*S*I/N - rate*I1
      dI2 = rate*I1 - rate*I2
      dI =  dI1 + dI2
      dR = rate*I2
      list(c(dS, dI, dI1, dI2, dR))
  })
}
```

With the transitions identified, user can then define the model in `denim`.

When using `denim` DSL, the model structure is given as a set of *key-value* pairs where

- *key* shows the transition direction between compartments in the format of `compartment -> out_compartment`.
- *value* is either a built-in distribution function that describe the transition or a mathematical expression.

```
# --- Model definition in denim
transitions <- denim_dsl({
  S -> I = beta * S * I/N
  # shape is 2 from number of I sub compartments
  I -> R = d_gamma(rate = 1/3, shape = 2)
})
```

#### Model configurations

Similar to `deSolve`, `denim` also ask users to provide the initial values and any additional parameters in the form of named vectors or named list.

For the example `deSolve` code, while users can use the `initalValues` from the `deSolve` code as is (`denim` will ignore unused `I1`, `I2` compartments as these sub-compartments will be automatically computed internally), it is recommended to remove redundant compartments (in this example, `I1` and `I2`).

For parameters, since `rate` is already defined in the distribution functions, users only need to keep `beta` and `N` from the initial parameters vector. We do not need to specify the value for `timeStep` variable as this is a special variable in denim and will be defined later on.

```
# remove I1, I2 compartments
denim_initialValues <- c(S = 999, I = 1, R=0)
denim_parameters <- c(beta = 0.3, N = 1000)
```

***Initialization of sub-compartments***: when there are multiple sub-compartments (e.g., compartment `I` consist of `I1` and `I2` sub-compartments), the initial population is always assigned to the first sub-compartment. In our example, since `I = 1`, denim will assign `I1 = 1` and `I2 = 0`.

There is also an option to distribute initial value across sub-compartments based on the specified distribution. To do this, simply set `dist_init` parameter of distribution function to `TRUE`.

```
transitions <- denim_dsl({
  S -> I = beta * S * I/N
  I -> R = d_gamma(rate = 1/3, shape = 2, dist_init = TRUE)
})
```

However, for comparison purposes, we will keep this option `FALSE` for the remaining of this demonstration.

#### Simulation

Lastly, users need to define the simulation duration and time step for `denim` to run. Unlike `deSolve` which takes a time sequence, `denim` only require the simulation duration and time step.

Since `denim` uses a discrete time approach, time step must be set to a small value for the result to closely follow that of `deSolve` (in this example, 0.01).

```
mod <- sim(transitions = transitions,
             initialValues = denim_initialValues,
             parameters = denim_parameters,
             simulationDuration = 100,
             timeStep = 0.01)
```

#### Compare output

The following plots show output from `denim` and `deSolve`

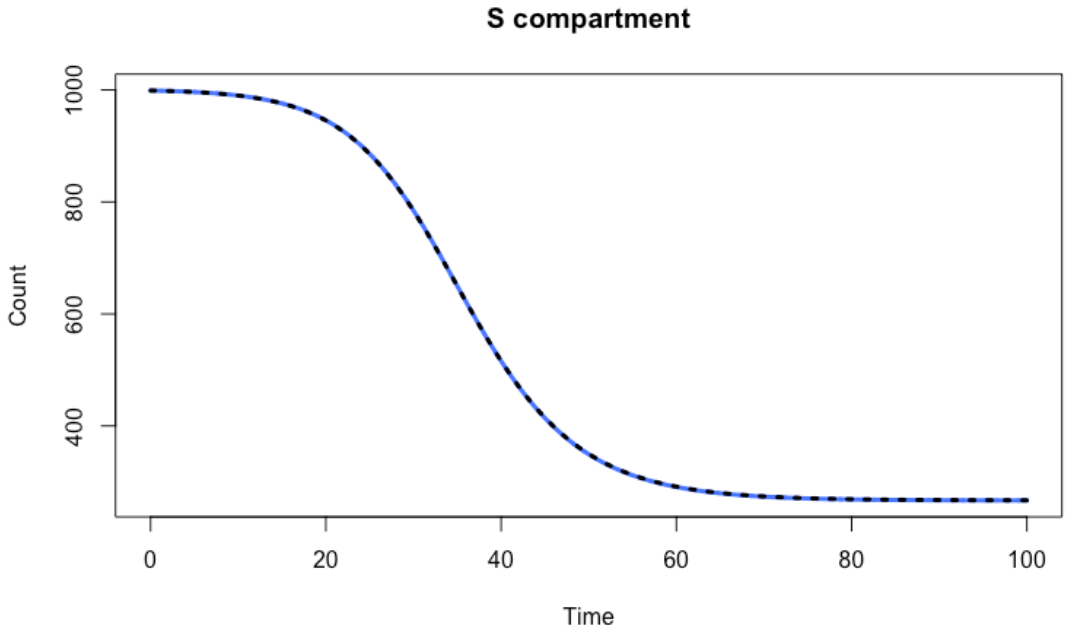

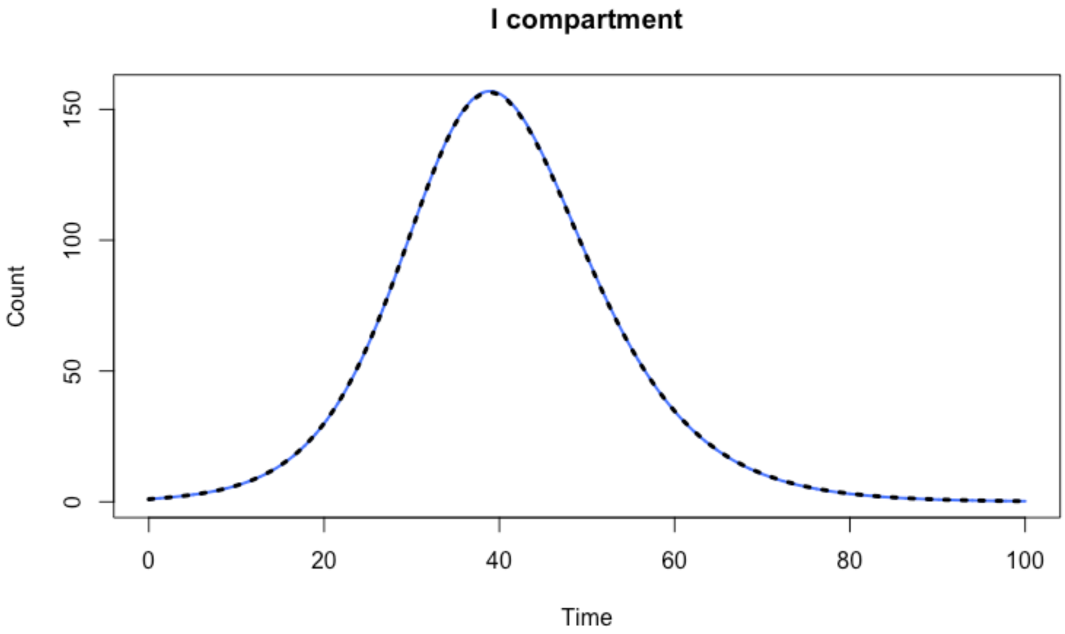
I compartment
Count
0
50
100
150
0
20
40
60
80
100
Time

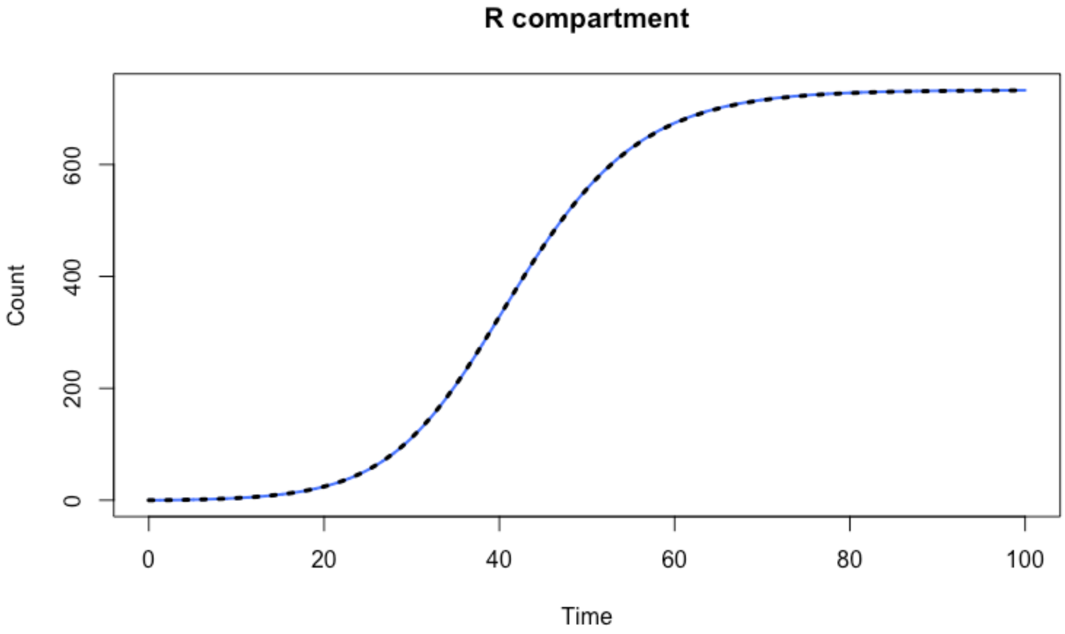
R compartment
Count
0
200
400
600
0
20
40
60
80
100
Time
